# Residual feed intake in dairy ewes: an evidence of intraflock variability

**DOI:** 10.1101/723809

**Authors:** E. González-García, J. P. Dos Santos, P. Hassoun

## Abstract

This study examined the intraflock variability of feed efficiency in dairy ewes, through monitoring residual feed intakes (RFI). Primiparous lactating ewes (*n*=43; 57.7±0.91 kg body weight [BW] at lambing), representative of a French Lacaune dairy flock, were allocated in an equilibrated 2 × 2 factorial design experiment, lasting for 63 days during mid-lactation and combining 2 litter sizes (singletons, SING or twins, TWIN) and 2 daily milking frequencies (once, ONE or twice, TWO). Ewes were individually fed a diet based on ryegrass silage, local hay and supplements. Individual DMI was recorded daily and further used to evaluate (and compare) differences in RFI between ewes at 35, 42, 49, 56, 63, 70, 77, 84, 91 and 98 days relative to lambing (DIM). Total (BW) and metabolic (BW^0.75^) body weight, BCS, milk yield and plasma NEFA were monitored weekly. Differences in DMI were mainly due to the lactation stage and litter size and were 11% higher in ewes with TWIN compared to SING. This was positively correlated to milk yield and consistent with differences in RFI which varied due to litter size and to the milking frequency × lactation stage interaction. Ewes that lambed SING showed higher feed efficiency (□0.13±0.020 vs. 0.08±0.015 kg DM/ewe/d of RFI in SING vs. TWIN, respectively), whereas there was no differences in BW or BCS. Milking frequency did not affect DMI but milk yields were higher in TWO, which was related to a higher feed efficiency in this group (0.04±0.017 vs. □0.10±0.018 kg DM/ewe/d of RFI in ONE vs. TWO, respectively). Average RFI was affected (*P* <0.0001) by the ewe, thus allowing a ranking among individuals to be established. High (n=22) or low (n=21) feed efficiency ewes averaged □0.17±0.09 or 0.18±0.09 kg DM/d RFI, respectively. Estimates of RFI were not correlated to the individual milk production potential. Even if no differences in BW, BW^0.75^ or BCS were detected, high efficiency ewes mobilised almost two-fold their body reserves when compared to the low efficiency group. The observed intraflock variability in feed efficiency of this dairy ewes flock was affected by litter size and milking frequency but also by evident differences between individuals’ physiologies.

## Introduction

It is well known that feed accounts for most of the total farm expenses in animal production systems and that a possible solution to lessen overall feed costs and alleviate the associated negative environmental impacts is to select for feed efficiency traits. In the past, producers have primarily focused on feed conversion ratios; however, animals with similar ratios differ in feed intake and productivity. As an alternative, Koch *et al*. (1963) proposed selecting residual feed intake (RFI), sometimes referred to as net feed intake. Considered the deviation of actual intake from the predicted intake for a given measure of growth (ADG) and body weight, RFI can be used to compare individuals with the same or differing levels of production during the period of measurement.

In contrast to feed conversion, selection based on RFI seems to select for lower rates of consumption and animal maintenance requirements than contemporaries to yield the same amount of product without changing adult weight or rate of gain, so theoretically these animals should cost less to feed on a daily basis when the costs of all other maintenance factors for these animals (breeding, health, etc.) are held constant (Bezerra et al., 2013; Potts et al., 2015). Heritability of RFI has been reported to be moderate, e.g. 0.06 to 0.24 depending on lactation stage (Tempelman et al., 2015) in dairy cattle e.g. 0.32□0.33 (Gonzalez-Recio et al., 2014; Veerkamp et al., 1995). Nevertheless, five major physiological processes are likely to contribute to variations in RFI, with these processes being associated with the intake and digestion of feed, metabolism (anabolism and catabolism associated with and including variation in body composition), physical activity, and thermoregulation (Herd and Arthur, 2009).

Thus, there is growing interest among producers with respect to using RFI as a tool for genetic improvement, with a greater experience in swine (Patience et al., 2015) and poultry; (Aggrey and Rekaya, 2013), numerous research efforts have investigated the effectiveness of selecting for feed efficiency using RFI in beef cattle (Fitzsimons et al., 2014; Gomes et al., 2012), dairy cattle (Green et al., 2013; Potts et al., 2015; Pryce et al., 2014) or sheep (Cockrum et al., 2013; Meyer et al., 2015; Redden et al., 2013). The sheep industry, however, has yet to fully investigate the potential impacts associated with selecting for RFI on carcass merit, growth traits, reproduction traits, and fleece characteristics (Cockrum et al., 2013). In the particular case of the dairy sheep sector, to our knowledge, there is no available information on RFI. Beyond its economic attractiveness, evaluating factors affecting intraflock variability in feed efficiency, also contributes to increasing our knowledge regarding the available spectrum of adaptive capacities that can be found at the intraflock level, when the interpretation of individual RFI is combined with other physiological processes like body reserves mobilisation-accretion.

The objective of this study was to evaluate factors affecting the intraflock variability in the feed efficiency of individually fed primiparous Lacaune dairy ewes, by focusing in the analysis of individual differences in RFI during several weeks in mid-lactation. We evaluated the hypothesis that variability in feed efficiency of individuals belonging to the same breed, cohort, productive purpose, with similar age, and reared under identical conditions, could be due to non-genetic factors but also to differences between the individuals. A second hypothesis was that mechanisms responsible for differences in RFI among individuals would probably be related to those linked to the use of body reserves.

## Materials and methods

### 2.1. Animals, management, treatments, and measurements

The experiment was carried out with a representative flock of primiparous Lacaune dairy ewes belonging to the INRA Experimental Farm La Fage, Causse du Larzac (43°54′54.52″N; 3°05′38.11″E; ~800 m above sea level), Aveyron, France, following the procedures approved by the Regional Ethics Committee on Animal Experimentation, Languedoc-Roussillon (France), Agreement 752056/00.

A detailed description of the animals, management and experimental design used for collecting the data employed in this study can be obtained from González-García et al. (2015). We used the data belonging to the primiparous (PRIM) group, considering that they were individually fed. Briefly, forty-three Lacaune dairy ewes (PRIM; two-tooth ewes) were chosen from the main flock at the end of pregnancy. Lambing took place in January (mean lambing date was 12 January) with a mean body weight (BW) of 57.7±0.91 kg. Litter size was determined about 2 months before lambing, by obtaining an ultrasound image for each ewe in the flock. The effects of two contrasting daily milking frequencies (once, ONE vs. twice, TWO) were evaluated. Approximately one half of the experimental flock was submitted to the ONE milking regime whereas their performance was compared to the other group, which was submitted to the conventional milking frequency (TWO).

The experimental ewes were thus distributed in homogeneous groups according to BW, BCS, and litter size and were allocated to a 2 × 2 factorial design according to litter size (lambing singletons [SING], n = 16 or twins [TWIN], n = 32) and daily milking frequency (FREQ; ONE, n=24; or TWO, n=24). Thus, the design finally comprised 4 randomly assigned balanced groups for which the mean BW at lambing was as follows: (1) SING×ONE (n=8; 57.2±2.46 kg); (2) SING×TWO (n=8; 59.7±2.47 kg BW); (3) TWIN×ONE (n= 16; 58.5±1.62 kg BW); (4) TWIN×TWO (n=16; 56.1±1.44 kg).

Ewes were housed in confinement in straw-bedded pens and had access to an individual feeding post controlled by an electronic device that allowed each animal to get into its correct place using individual electronic identification (IDE). Each ewe-lamb was equipped with an IDE ear tag that recorded its presence at the feed bunk and allowed (or not) access to the individual feeder. When a ewe approached the feed bin, the unique passive ear transponder was identified, the barrier was unlocked, and the animal was allowed access to the feed.

Ewes were thus individually fed with a standard *ad libitum* lactating total mixed ration for dairy ewes composed of a 55% dry matter (DM) ryegrass silage, 18% hay (28% second cut alfalfa-cocksfoot and 72% of a third harvest local hay called *Foin de Crau*, composed of a multiple mixture of grasses, legumes, and other species), 13% barley grain, 9% dehydrated alfalfa (*Luzapro*, 26.5% crude protein), and 6% commercial concentrate (*Brebitane*, 46% crude protein). The total mixed ration was offered twice daily, one-third in the morning and two-thirds in the afternoon, at about 9 AM and 6 PM, respectively. Its distribution was adjusted to an allowance rate of 115% of the previous day’s voluntary intake. In addition, 90 g DM of *Brebitane* was offered at each milking in the milking parlour, or twice this amount in the morning for the group that was milked once a day. Ewes had free and continuous access to fresh water and salt block.

### 2.2 Determination of residual feed intake (RFI) and monitoring related zootechnical and metabolic parameters

The quantities of feed offered and refused were recorded daily in order to determine the individual daily actual feed intake and thus the daily dry matter intake (DMI) per ewe. Average DMI was thus individually calculated weekly, and further used to evaluate (and compare) individual RFI per ewe. Expected feed intake was calculated based on the INRA table recommendations for dairy sheep, according to their lactation stage, BW and overall physiological requirements (Hassoun and Bocquier, 2014). The RFI was calculated as the difference between the expected feed intake (Hassoun and Bocquier, 2014) and the actual, individually measured feed intake in the experiment.

Measurements of RFI were scheduled at 35, 42, 49, 56, 63, 70, 77, 84, 91 and 98 days relative to lambing (or days in lactation, DIM). Around each sampling date, ewes were individually weighed, BCS was assessed by trained observers using the 5-point scale proposed by Russel et al. (1969) and a blood sample was taken before the first meal distribution for metabolic profile (i.e. plasma non-esterified fatty acids, NEFA; González-García et al., 2015). Here, we use data referring to the plasma concentration of NEFA, as an indicator of body reserves mobilisation.

Ewes were milked twice daily at 8 AM and 5 PM. Machine milking was performed in a double-24 stall parallel milking parlour. Milk yield and milk composition (fat and protein content) were monitored and standardised milk yield (SMY) calculated (Bocquier et al., 1993).

### 2.3 Data processing and statistical analyses

In the first step, the effects of major sources of variation i.e. lactation stage (35□98 DIM), litter size, milk frequency, and their first-order interactions on the main variables of interest i.e. RFI (kg/ewe/day) and other zootechnical parameters (i.e. dry matter intake –DMI-, RFI, and actual or standardised milk yields –SMY-), linked to the feed efficiency of these primiparous Lacaune dairy ewes, were determined by using the PROC MIXED procedure of SAS (SAS; v. 9.1.3., 2002□2003 by SAS Institute Inc., Cary, NC, USA) with the following statistical model:

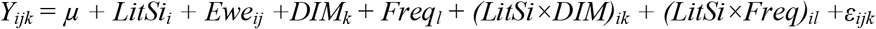

where *Y*_*ijk*_ is the response at time *k* on ewe *j* with litter size *i*, *µ* is the overall mean, *LitSi*_*i*_ is a fixed effect of litter size *i* (_*i*_ = 1–2), *Ewe*_*ij*_ is a random effect of ewe *j* with litter size *i*, *DIM*_*k*_ is a fixed effect of time or days relative to lambing (DIM; 35□98) *k*, *Freq*_*l*_ is a fixed effect of daily milking frequency *l* (_*l*_ = 1–2), *(LitSi×DIM)*_*ik*_ is a fixed interaction effect of litter size *i* with time *k*, *(LitSi×Freq)*_*il*_ is a fixed interaction effect of litter size *i* with daily milking frequency *l* and *ε*_*ijk*_ is random error at time *k* on ewe *j* with litter size *i*.

In a second step, and after the determination of RFI, ewes were classified into high or low feed efficiency individuals based on their distribution in this experimental population, when considering their average RFI values determined for the whole experimental period (i.e. from 35 to 98 DIM). The analysis of variance (developed in the first step), allowed the level of variation at each significant intra-factor level to be analysed in detail with regard to the main variable of interest i.e. feed efficiency through RFI. Using the PROC RANK of SAS, the average ranking of the individual ewes for the RFI variable was established. The same procedure allowed the experimental ewes to be classified as high, medium and low milk producers. The last allowed a relationship between the individual RFI and the SMY potential of each ewe to be established using the PROC REG of SAS. The dependency of feed efficiency from milk yield potential was thus analysed.

In the final step, once the ewes were classified as tending to belong to the high or low feed efficiency groups, the relationships between the RFI and the average total (BW) or metabolic body weight (BW^.75^), BCS and plasma NEFA profile were evaluated using the PROC GLM procedure of SAS. The general statistical model used for this was as follows:

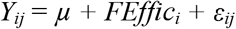

where *Y*_*ij*_ is the observation, *µ*, the population mean, *FEffic*_*i*_, the feed efficiency rank effect (_*i*_ = 1–2; low or high) and *ε*_*ij*_ is the residual error.

For all traits, the experimental unit was considered the ewe, as they were individually fed and included in the model as a random effect. Significance was declared at probability levels of ≤5% and comparisons between means were tested with the least squares means (LSMeans) separation procedure using the PDIFF option of SAS.

## Results

The statistical significance of the lactation stage, litter size, milking frequency and first-order interactions on DMI, milk yield and RFI are presented in Table 1. Observed differences in DMI were mainly due to the lactation stage (DIM) and litter size effects, but not because of changes in milking frequency *per se*. The effects of milking frequency on DMI depended on its interaction with litter size. Similarly to milk yields, RFI was strongly affected (*P* <0.0001) by the three major sources of variation evaluated here (i.e. DIM, litter size and milking frequency), and by the interaction milking frequency × lactation stage. A similar tendency was observed for SMY (Table 1).

**Table 1.** Significance (*P* value) of the fixed effects lactation stage (days in milk, DIM), litter size (LS), milking frequency (MF) and their interactions on residual feed intake (RFI, kg/ewe/day) and other zootechnical parameters linked to feed efficiency of primiparous Lacaune dairy ewes during lactation (35□98 DIM).

|  | DIM | Litter size | Milking frequency | First-order interactions across major fixed effects |  |  |  |
| --- | --- | --- | --- | --- | --- | --- | --- |
|  |  |  |  | LS×MF | LS×DIM | MF×DIM | LS×MF×DIM |
| DMI | 0.0165 | <.0001 | 0.2694 | 0.0030 | 0.9926 | 0.9748 | 0.8628 |
| MY | <.0001 | <.0001 | <.0001 | 0.0428 | 0.9880 | 0.2395 | 0.9814 |
| SMY | <.0001 | 0.0105 | <.0001 | 0.0052 | 0.9837 | 0.0187 | 0.9413 |
| RFI | <.0001 | <.0001 | <.0001 | 0.3539 | 0.7881 | <.0001 | 0.2485 |

Average DMI during the evaluated mid-lactation period was 11% higher in ewes that lambed twins when compared to those lambing singletons, and was positively correlated (data not shown) to the total or SMY milk yields (Table 2). Differences (*P* <0.0001) in RFI were also found between ewes that lambed SING and TWIN (□0.13±0.020 *vs*. 0.08±0.015, respectively; Table 2). Thus, despite a lower milk yield, ewes that lambed singletons converted feeds more efficiently.

**Table 2.** Effects of litter size and milking frequency on dry matter intake (DMI, kg/ewe/day), actual or standardised (SMY) milk yields (l/ewe/day) and residual feed intake (RFI, kg/ewe/day) of individually fed primiparous Lacaune dairy ewes at mid-lactation (35□98 DIM).

| Item | Litter size |  | Milking frequency |  |
| --- | --- | --- | --- | --- |
|  | SING | TWIN | ONE | TWO |
| Dry matter intake (DMI, kg/ewe/day) | 2.14±0.028 | 2.38±0.021 | 2.28±0.024 | 2.24±0.025 |
| Milk yield (kg/ewe/day) | 1.59±0.032 | 1.77±0.024 | 1.57±0.028 | 1.80±0.029 |
| Standardised milk (SMY, L/ewe/day) | 1.37±0.024 | 1.45±0.018 | 1.32±0.021 | 1.50±0.022 |
| RFI (kg DM/ewe/day) | -0.13±0.020 | 0.08±0.015 | 0.04±0.017 | -0.10±0.018 |

Milking frequency did not affect DMI (Table 1) but, as expected, the actual or SMY milk yields were higher in ewes being milked twice (Table 2) which was related to a higher overall feed efficiency in this group for the whole experimental period, as interpreted by differences in RFI (0.04±0.017 vs. □0.10±0.018 in ewes milked once *vs*. twice a day, respectively; Table 2).

Those effects of milking frequency on feed efficiency were dependent on the lactation week (DIM; Figure 1) (see also significant interactions between milking frequency and DIM on RFI; Table 1). Only at 35 DIM, RFI was lower in ewes milked once daily when compared to those milked twice. This tendency changed from the second week (42 DIM) until the end of the experiment (98 DIM), the period during which ewes belonging to the group TWO were always more efficient, as interpreted by their lower RFI, with regard to ewes from the group ONE (Figure 1).

**Figure 1.**
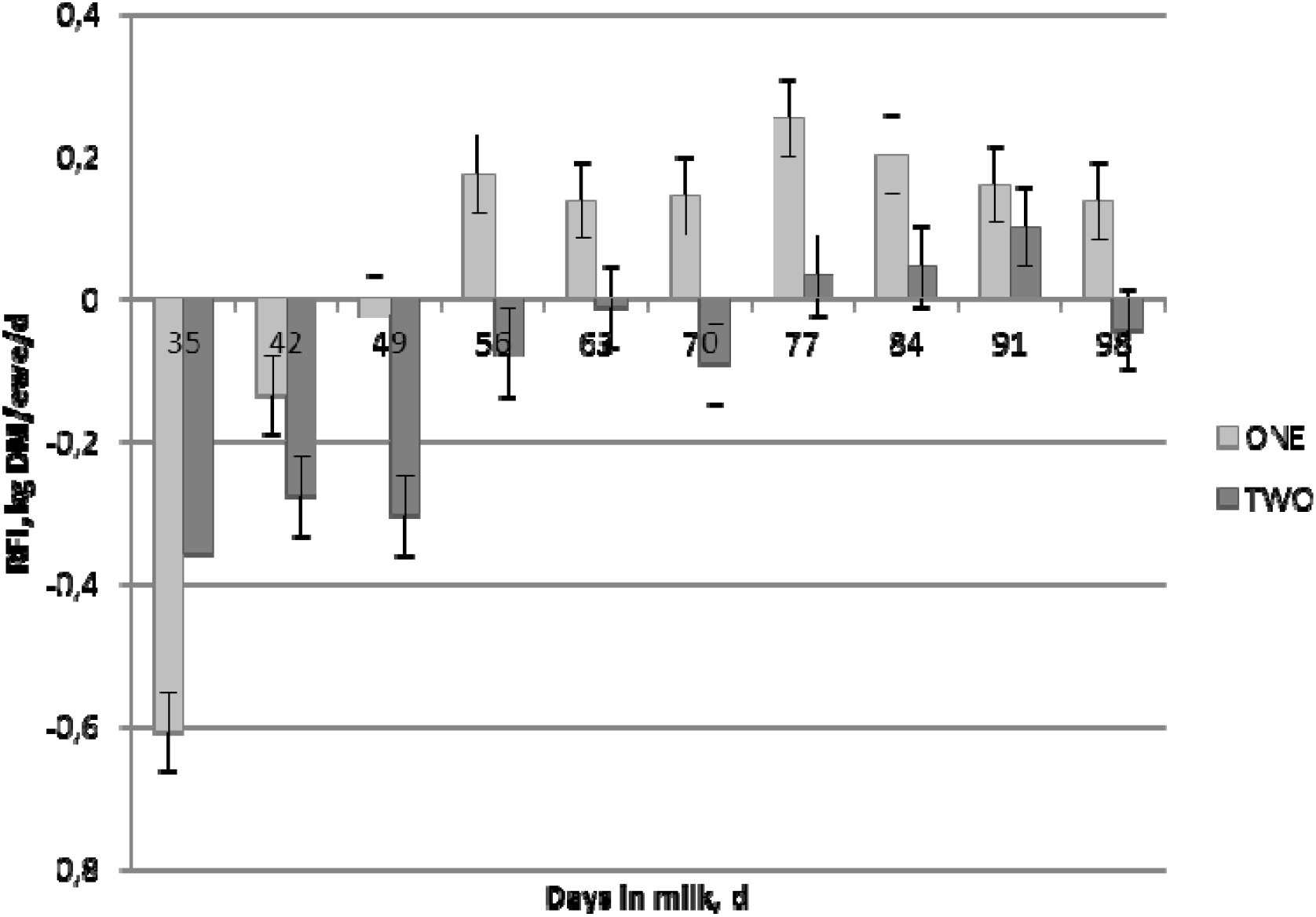
Effects of the lactation stage (days in milk, DIM) and its interaction with milk frequency on residual feed intake (RFI) of primiparous Lacaune dairy ewes during mid-lactation (35□98 DIM).

The average RFI for the whole experimental period was significantly (*P* <0.0001) affected by the ewe itself. As a consequence, ewes were ranked as having a tendency to high or low feed efficiency as a function of their average RFI (Figure 2). Ewes classified as high feed efficiency ewes (n=22) averaged □0.17±0.09 kg DM/d of RFI; whereas, on the other hand, ewes classified as low feed efficiency ewes (n=21) showed an average RFI value of 0.18±0.09 kg DM/d.

**Figure 2.**
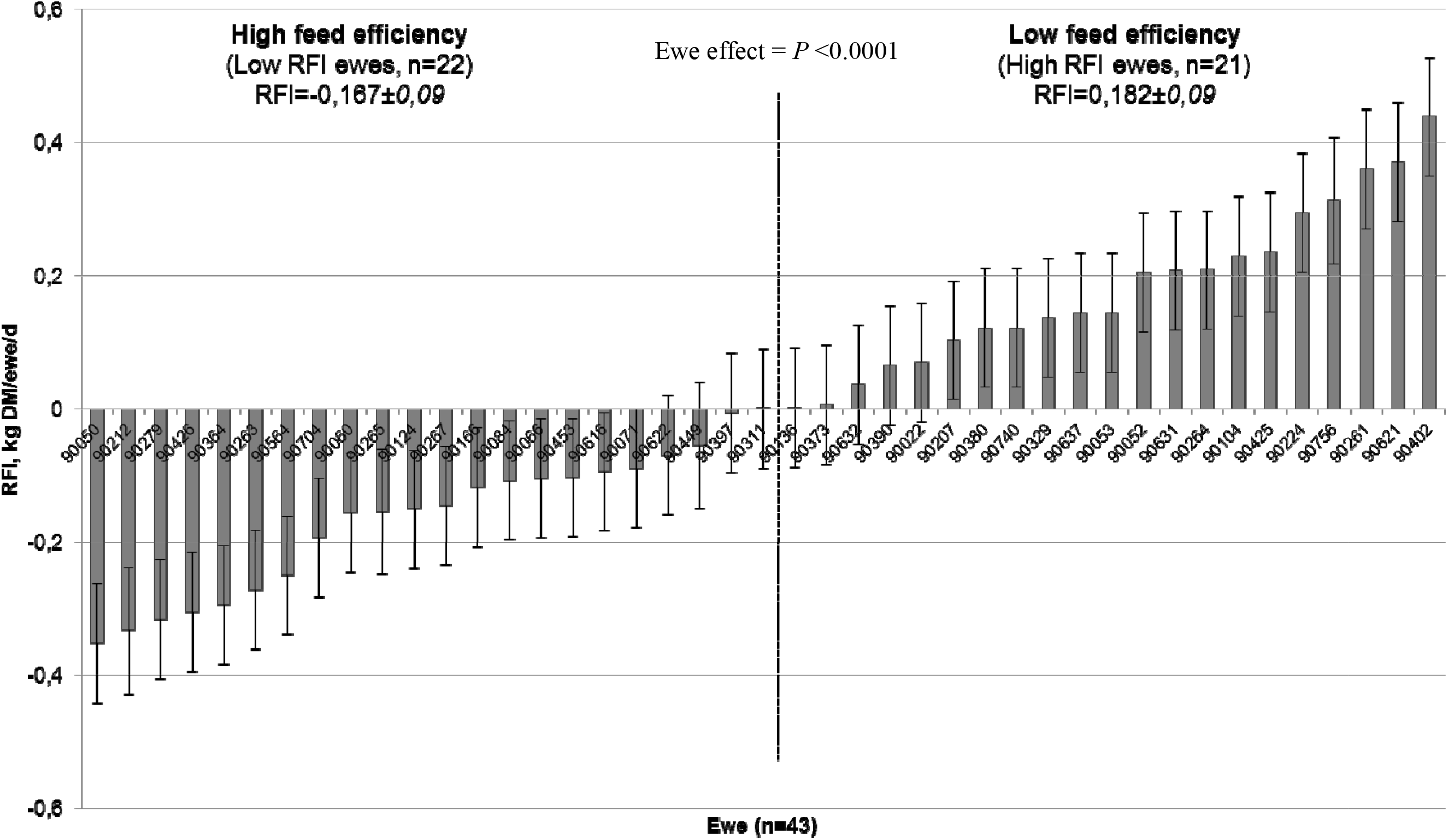
Ranking of primiparous Lacaune dairy ewes (n=43) in function of their average residual feed intake (RFI, kg DM/ewe/day) during mid-lactation (35-98 days in milk). Overall RFI during the whole period was 0.004±0.090 kg DM/ewe/d.

The expected RFI was independent of the individual milk production potential (Figure 3). In more than half of the cases, ewes classified as tending to be high feed efficiency ewes (left side panel of Figure 2) corresponded to ewes submitted to two milking per day (13 ewes in TWO and 9 in ONE *vs*. 6 in TWO and 15 in ONE in high vs. low efficiency groups, respectively; Table 3).

**Table 3.** Relationships between residual feed intake (RFI) and average total (BW) or metabolic body weight (BW^.75^), body condition score (BCS) and plasma non-esterified fatty acids (NEFA) profile in individually fed primiparous Lacaune dairy ewes at mid-lactation (35□98 DIM). Ewes were classified as presenting high or low feed efficiency in accordance to their average RFI during the 8 week period. NS = non-significant; LS = litter size; MF = milking frequency.

| Item | Feed efficiency of ewes, according to RFI ranking |  |  |  | Effect, *P*<br>value |
| --- | --- | --- | --- | --- | --- |
|  | High |  | Low |  |  |
| | LSmean | S.E.M. (±) | LSmean | S.E.M. | |
| BW, kg | 59.0 | 1.46 | 56.3 | 2.81 | NS |
| BW<sup>0.75</sup>, kg | 21.3 | 0.39 | 20.5 | 0.99 | NS |
| BCS, 1–5 | 2.88 | 0.035 | 2.77 | 0.132 | NS |
| NEFA, $\mu\text{mol/L}$ | 0.39 | 0.017 | 0.22 | 0.016 | <.0001 |
| LS distribution | 13 SING; 9 TWIN |  | 3 SING; 18 TWIN |  |  |
| MF distribution | 9 ONE; 13 TWO |  | 15 ONE; 6 TWO |  |  |

**Figure 3.**
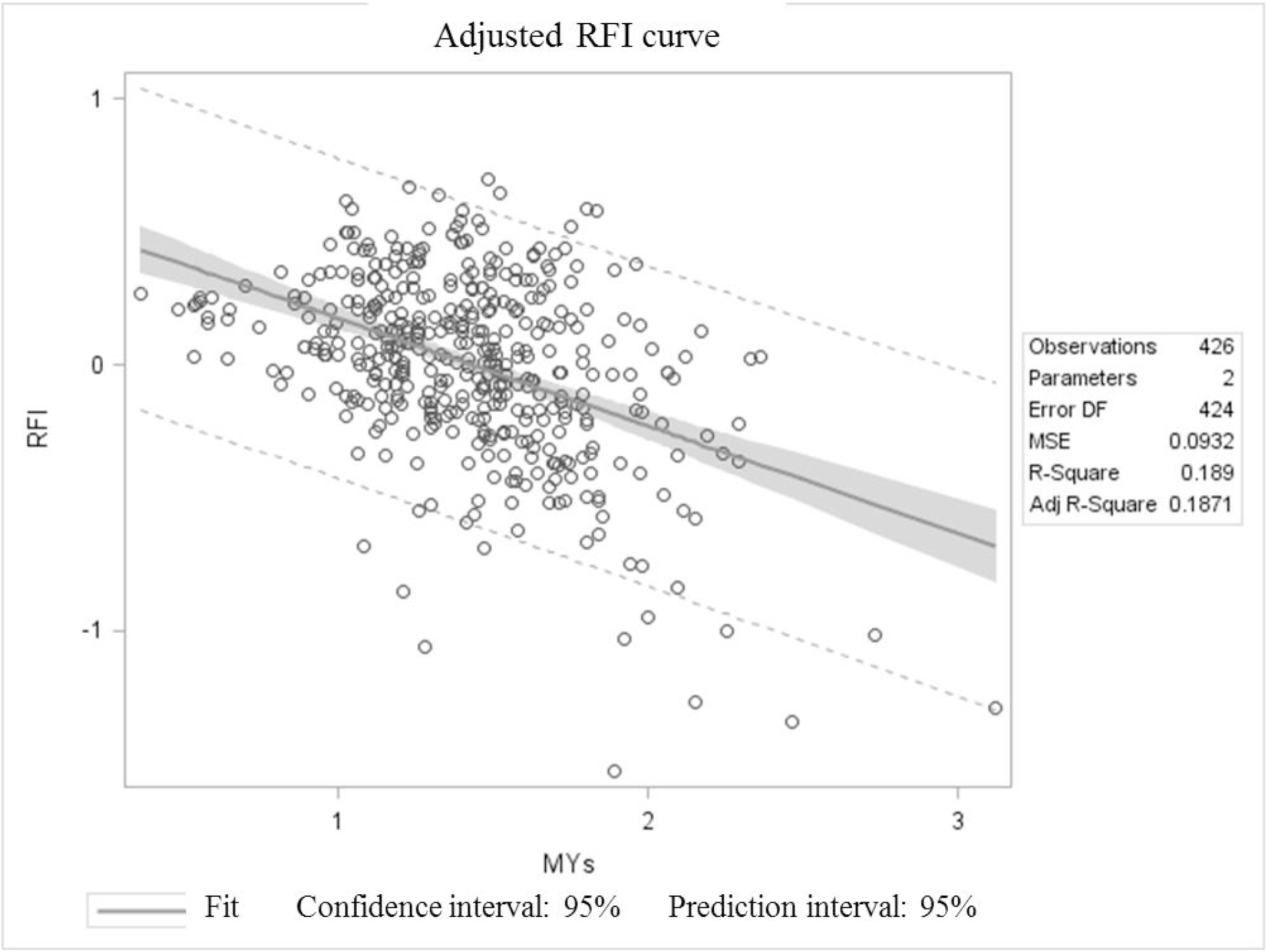
Adjusted curve for average individual residual feed intake (RFI) and fat-corrected milk yield of primiparous Lacaune dairy ewes (n=43) during mid-lactation (35-98 days in milk). MYs= standardise milk yield (SMY).

Interestingly, and even if no differences in BW, BW^0.75^ or BCS were detected, high efficiency ewes mobilised almost two-fold their body reserves when compared to the low efficiency group (see and compare plasma NEFA values in Table 3). The latter probably supported a higher energy requirement for milk production, considering the larger proportion of ewes being milked twice (TWO) in the high efficiency group. However, three of the four most efficient ewes that lambed singletons and were milked once a day, which illustrate the fact that intraflock variability in feed efficiency is also affected by differences in individual natures among animals belonging to the same breed, cohort and receiving the same management, and their implicit, not well known related mechanisms.

## Discussion

Evaluating factors affecting intraflock variability of feed efficiency, through RFI, increases our knowledge regarding the available spectrum of adaptive capacities which can be found at the intraflock level; this becomes more interesting when the interpretation of RFI is combined with other physiological processes like body reserves mobilisation-accretion. However, the exercise is also interesting from an economic point of view for the industry in question since the identification of animals that require less feed for normal production would clearly increase overall farm productivity, thus leading to the argument that feed conversion efficiency of farm animals could be considered an important component of the profitability of farming systems (Cockrum et al., 2013; Pryce et al., 2014; Redden et al., 2013; Williams et al., 2011).

The RFI quantifies efficiency within a production level and allows for the identification of animals that convert gross energy into net energy more efficiently by reducing energetic losses in faeces, urine, gas, and nonmaintenance heat; thus, this is independent of the dilution of maintenance when multiple of maintenance is calculated based on requirements for observed production (Potts et al., 2015).

Currently, there are several reports arguing the interest, pertinence and possibilities of using RFI as a selection characteristic to increase feed efficiency and farm profitability in non-ruminants (Patience et al., 2015), but also in ruminants (beef: Fitzsimons et al., 2014; Gomes et al., 2012; dairy: Green et al., 2013; Potts et al., 2015; Pryce et al., 2014). There is a lack of information, however, in small ruminants, although some works have been developed mainly during the growth phase in sheep (Cockrum et al., 2013; Meyer et al., 2015; Redden et al., 2013) and the sheep industry has yet to fully investigate the potential impacts associated with selecting for RFI on carcass merit, growth traits, reproduction traits, and fleece characteristics (Cockrum et al., 2013).

In the dairy sheep industry, to our knowledge there is no available information on RFI studies. Our work and results could thus be considered original in that sense. The question of using RFI as a tool for increasing feed efficiency in the future dairy ewe’ industry remains.

Here, we evaluated different factors with the potential for affecting feed intake i.e. litter size at lambing and during the suckling period, milking frequency and lactation stage. However, we were also able to confirm evidence for individually intrinsic factors leading to differences in feed efficiencies at the intraflock level in ewes belonging to the same dairy breed, cohort, with the same age and reared under identical conditions, using the same day to day management and feeding.

Differences between the energy requirements of ONE and TWO milking frequencies and ewes lambing SING or TWIN were great enough to cause significant differences in energy partitioning, which were translated into differences in milk yield and feed intake. Thus, our findings provide support for the use of RFI as a tool to identify animals that eat less than others within a production level, independent of the energy balance or the related management practice.

Although some re-ranking occurred, this was minor, so that within a level of production, ewes that eat less than their contemporaries when receiving a particular management (e.g. milked once daily) should also consume less than their contemporaries when returning to the average management of the flock (e.g. being milked twice a day).

We also verified that RFI was independent of the individual milk production potential (Figure 3). Thus, we could assume that ewes with low RFI required less feed to produce the same amount of product as their contemporaries, independently of their milk production potential. Consistent with these results, the most feed-efficient ewes (n = 22) ate ~350 g of DM/d less than the least efficient ewes in our study (i.e. □0.17±0.09 *vs*. 0.18±0.09 kg DM/d of RFI in high and low feed efficiency ewes, respectively).

Potts et al. (2015) reported that RFI was moderately repeatable across 2 consecutive feeding periods for replacement beef heifers classified as high (>0.5 SD), medium (±0.5 SD), and low (<−0.5 SD) RFI. These authors argued that the estimation of RFI across different periods may be more repeatable if measurements are obtained from periods when animals were in similar physiological states. There is little to no information available on those effects in cattle or sheep, but species may differ in their response to RFI selection (Cockrum et al. (2013).

Here, we focused on the mid-lactation of this Lacaune dairy flock, an important period on which a relative stabilisation is achieved; thus, comparison among individuals becomes feasible. The weekly estimates of RFI were also repeatable within and across group of ewes in the design, which suggests that we were able to account for many of the production and BW changes that occurred from one week to the next, affecting the calculation of RFI. However, genotype × environment interactions may be an important factor to take into account.

While upwards of 60 d of feed intake measurements are needed to accurately estimate RFI in beef cattle (Sainz and Paulino, 2004), the necessary duration in sheep is unknown (Cockrum et al., 2013). In our study, we chose a period (63 days) similar to that recommended by Sainz and Paulino (2004) which also fits well in the range indicated by Cockrum et al. (2013) (42–63d) for determining RFI in sheep. The last authors confirmed that both the variance and the R² of their results in rams provide support that a period of 40–63 d is needed to accurately determine individual RFI values in sheep.

The current study contributes to the literature on the relationship between RFI and productive performance in dairy ewes offered a forage diet. To date, the majority of studies examining this area in sheep have focused on growing or finishing animals offered energy-dense diets.

Similar to reports in beef cattle, our RFI results varied widely in dairy ewes (see standard errors in Figure 2). This variation may be attributed to individual differences in heat production from metabolic processes, body composition, and physical activity; these factors account for around 70% of the variation in RFI (Herd and Arthur, 2009), but were not measured in this study.

Results indicated that BW, BCS and milk production potential are phenotypically independent of RFI estimates but further research is necessary to determine the relative weighting value for RFI in successive physiological states before it can successfully be considered a potential selection tool.

Based on their results, Cockrum et al. (2013) recommended that, ideally, selection decisions for RFI should take place at weaning, and feed efficiency status should be applicable over an animal’s lifetime.

Fitzsimons et al. (2014) found changes in backfat thickness, which were negative for low-RFI cows and positive for high-RFI cows. The reduction in backfat thickness in low-RFI cows suggested that these cows were mobilising more body fat to meet their nutritional requirements during pregnancy than high-RFI cows. These authors also suggested that the calculation of RFI in beef cows should include body composition traits such as subcutaneous body fat and BCS.

Potts et al. (2015) also argued that because body energy changes are accounted for in the prediction of RFI, it is expected that cows with low RFI will not be any more likely to mobilise body tissue to support production than cows with high RFI. The independence of RFI from BW loss is important because excessive tissue mobilisation can lead to negative energy balance, which is related to metabolic diseases and poor fertility.

Our results are in agreement with statements relating feed efficiency with body reserves utilisation. We found a positive correlation between RFI and the profile of body reserves mobilisation, analysed through regular monitoring of plasma NEFA, with the low-RFI ewes consistently showing higher plasma NEFA. Even if no differences in BW, BW^0.75^ or BCS were detected, high efficiency ewes mobilised almost two-fold their body reserves when compared to the low efficiency group. In the available literature, we did not find any previous report concerning the direct relationships between efficiency in body reserves administration and RFI in small ruminants and particularly in dairy sheep, as evidenced in the current study.

## Conclusions

Under the conditions of this experiment, low-RFI lactating dairy ewes ate less and produced similar levels when compared to high-RFI cohorts, without changing body weight or BCS; thus, they could be said to use their feed more efficiently. The observed intraflock variability in feed efficiency is probably the consequence of indirect factors affecting metabolism and energy balance of ewes, like litter size and milking frequency, but it is also affected by differences among the individuals’ natures. However, entry into different physiological stages may present some challenges and more research will be needed to investigate the long-term applicability of the RFI estimates found here. Finally, this is probably the first report demonstrating a close relationship between RFI and body reserves mobilisation in small ruminants, and particularly in dairy ewes.

## Acknowledgements

The authors are grateful to the technical staff of La Fage experimental farm unit for assisting with animal care, monitoring tasks and data collection.

